# Analysis of RAS as a tumor suppressor

**DOI:** 10.1101/153692

**Authors:** Chelsey Kenney, Edward C. Stites

## Abstract

The RAS GTPases are among the best-understood oncogenes that promote human cancer. Many have argued that non-mutated, wild-type, RAS also functions as a tumor suppressor. The arguments for RAS tumor suppressor activity often involve data that are claimed to be inconsistent with known principles of RAS biology. RAS tumor suppressor activity is invoked to explain these observations. Here, we consider an alternative hypothesis: these data are actually consistent with RAS biology. We investigate by using our previously developed mathematical model of RAS regulation. We find that three of five arguments for RAS having tumor suppressor activity are based upon data that the model demonstrates are actually consistent with known RAS biology. We also find that the other two types of data interpreted to indicate a tumor suppressor effect can be explained by our model with the additional assumption that RAS protein expression does not vary proportionally with gene dosage. Measurements of RAS protein expression as a function of gene dosage could help resolve whether or not RAS has tumor suppressor activity. Overall, we conclude that the evidence for RAS having tumor suppressor activity is much less strong that it has appeared.

## INTRODUCTION

Cancer is a disease where a subset of cells proliferates persistently to the detriment of the individual. The current theory states that cancer develops after cellular DNA acquires a collection of mutations that together provide all of the cellular behaviors needed for the ensemble cancer behavior. Mutations that promote cancer have been broadly divided into two classes that correspond to the patterns with which mutations occur. The first class is comprised of the oncogenes. Oncogenes are mutated genes that tend to promote increased activity compared to their original, non-mutated form (the protooncogene). The mutations that convert proto-oncogenes to oncogenes may disrupt the protein’s regulation to result in an oncogene with a higher propensity to promote the cancer-associated behavior. In general, a mutation to one of the two copies of a protooncogene is sufficient to induce the cancer promoting effects of the well-established oncogenes.

The second class is comprised of tumor suppressor genes. Tumor suppressor genes (TSGs) typically code for a protein with an action that counteracts a cancer promoting behavior. Mutations to tumor suppressor genes typically result in loss of function for the coded protein. This can be by deletion of the gene from the chromosome, by modifications to DNA or the chromosome that suppress gene transcription, or by mutations to the DNA that code for less functional protein, such as by a mutation that disrupts the protein’s activity. In general, both copies of a tumor suppressor gene are compromised in cancer because one copy is typically sufficient to maintain the tumor suppressive activity.

Among the most studied, most common, and best-understood oncogenes are the RAS oncogenes that occur within the *KRAS*, *NRAS*, and *HRAS* genes [1-3]. The RAS GTPases transmit cellular proliferation signals when they are bound to GTP, and the oncogenic mutations result in protein with a higher proportion bound to GTP. Several interesting experimental observations have been used to argue wild-type RAS can also behave as a tumor suppressor to counteract signals from oncogenic mutants [4-13] (Figure 1). Many of these experiments are difficult to mentally reconcile with known patterns of RAS signaling. However, biological networks are capable of many non-intuitive behaviors. Here, we consider the hypothesis that some of the data used to argue RAS has tumor suppressor activity are actually fully consistent with the oncogenic role of RAS. We evaluate the logical link between the experiments and known RAS biology by using a previously developed mathematical model of the processes that regulate the nucleotide binding state of RAS [14]. First, we evaluate five different types of experimental data that have been used to argue that RAS has tumor suppressor activity. We find that three types of data are consistent with our model of RAS regulation, demonstrating that new biology is not needed to explain these data. We find that the other two types of data are not consistent with the model, which suggests that processes not in the model, such as RAS having tumor suppressor activity, may be at play. However, we computationally demonstrate that it is not necessary to assume that RAS has tumor suppressor activity to explain these data. We do this by proposing an alternative hypothesis and then computationally demonstrating how this alternative hypothesis could also explain the data. We conclude that the argument that RAS has tumor suppressor activity may be less strong than it has appeared. We also describe specific experimental measurements that would test our alternative hypothesis.

**Figure 1:**
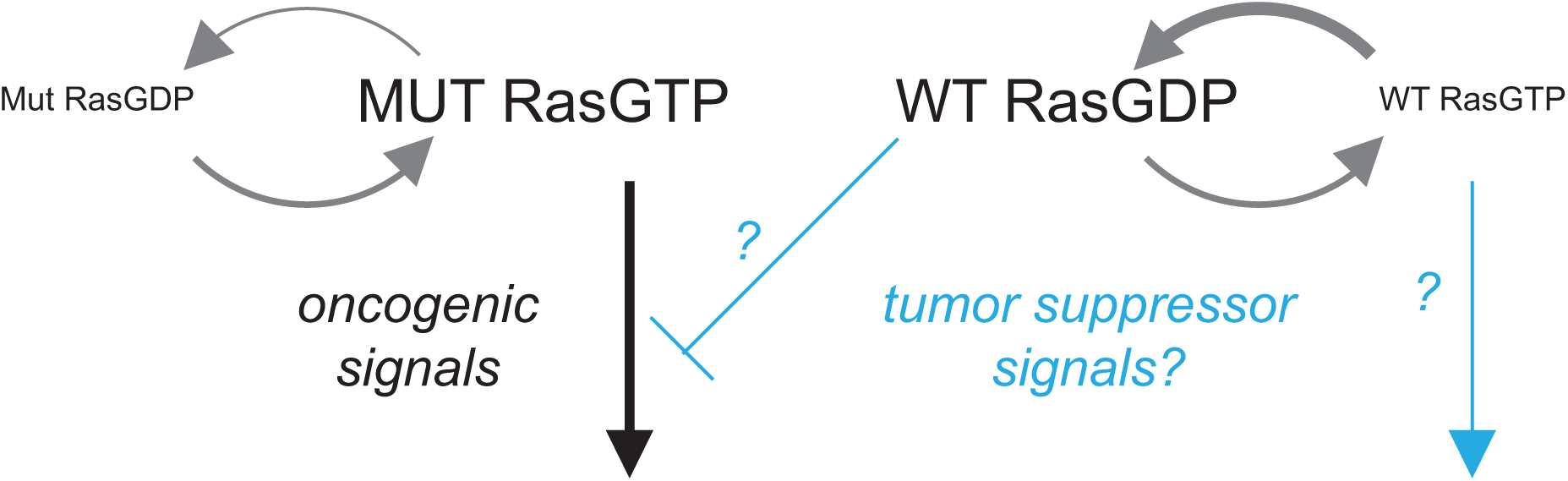
The RAS is a Tumor Suppressor hypothesis. Oncogenic RAS mutants are predominantly GTP bound and transmit proliferation signals. Wild-type RAS is predominantly GDP bound. It has been argued that wild-type RAS must transmit tumor suppressor signals that counteract oncogenic RAS signals to explain some experimental data. The tumor suppressor signal is indicated with questions mark to highlight that it is inferred, but not mechanistically understood.

## RAS signaling and the RAS signaling model

RAS signaling is largely dependent upon whether RAS is bound to GTP or GDP. Many proteins have RAS binding domains that specifically bind with RAS GTP. The interaction between RASGTP and a RAS binding domain protein typically contributes to the regulation of that RAS binding partner. Whether RAS is bound to GTP or GDP is influenced by several different processes [15]. RAS itself is a GTPase, which means that it can convert GTP to GDP. RAS GAPs (GTPase Activating Proteins) can also catalyze this reaction. Nucleotides can spontaneously dissociate and associate, providing another mechanism by which RAS can switch between nucleotide binding states. RAS GEFs (Guanine nucleotide Exchange Factors) can catalyze nucleotide exchange. RASGTP can further be sequestered by RAS effector proteins, which bind RASGTP through their RAS binding domain and thereby also prevent the bound RASGTP from interacting with GAPs or GEFs. The relative amount of RASGTP to RASGDP in the cell reflects the combined activity of all of these reactions.

We have previously developed and reported a computational model of RAS signaling that includes these different processes that regulate RAS signaling [14]. Massaction kinetics are used to model the reactions. The GAP and GEF reactions are modeled with irreversible and reversible Michaelis-Menten kinetics, which themselves are derived from mass-action kinetics. The parameters for these reactions are all based upon experimental measurements of these rate constants and enzymatic parameters. Simulations are used to find steady-state levels of RASGTP and RASGTP-effector complex. Parameters are also available to describe many of these same reactions for several oncogenic mutants. Among them is the mutation that replaces the Glycine (G) at codon 12 with an Aspartic Acid (D), also referred to as the RAS G12D mutation. The parameters that have previously been measured intuitively explain why the oncogenic mutants are more activating. For RAS G12D, the rate of GTP hydrolysis catalyzed by GAPs is essentially completely eliminated, resulting in a shifting of the dynamic equilibrium toward increased amounts of RASGTP. In this manner, our RAS model emergently results in increased patterns of RAS signal that are found with RAS pathway oncogenes [14]. The model also emergently results in increased patterns of RAS signal that are found with RAS pathway tumor suppressor genes [16]. Going forward, we use our model to investigate the experimental measurements that have been argued to support a tumor suppressor role for wild-type RAS. We use our same model and same parameters as previously published and described in detail [14, 17]. We assume simply that increased RAS signal results in increased cancer promoting potential, and that by varying variables of the model that correspond to biological variables that we can acquire insight into what happens within the biological system.

## ANALYSIS

### Analysis of Argument 1: Loss of Heterozygosity is common at the RAS allele

Loss of Heterozygosity (LOH), where one of the two distinct copies of a gene is lost and only one of the copies remains, is commonly observed for tumor suppressor genes. LOH may indicate a loss of one copy so that only one copy remains. Alternatively, it may indicate that one copy has been replaced with the other copy. LOH is commonly observed for tumor suppressor genes. LOH has been found to be common for *KRAS* [18]. When this was described, the authors also noted that several oncogenes (e.g. *HER2*) also display LOH so that LOH in itself is not sufficient to identify a TSG. In these early studies, it was also noted that LOH might indicate replacement of the wild-type allele with the mutant allele to increase mutant dosage [18]. Although frequent LOH for a gene is not uniquely indicative of it being a TSG, frequent LOH is occasionally cited as part of the larger argument for RAS acting as a TSG [19]. We evaluate the potential effects of dosage with our model.

We modeled the two possible situations that would display LOH (Figure 2A,B). To start, we considered 50% mutant and 50% wild type for the mutant/wild-type case (G12D/WT). We considered the apparent LOH situation where all of the wild-type allele is lost (G12D/-) by modeling this condition to have the same quantity of mutant G12D RAS protein as in G12D/WT conditions, but with all wild-type RAS removed from the model. We also considered the apparent LOH situation where the wild-type allele is replaced with the mutant allele (G12D/G12D). For the G12D/G12D conditions, we assumed that the concentration of the G12D protein was twice that used to model G12D/WT and G12D/-conditions.

**Figure 2:**
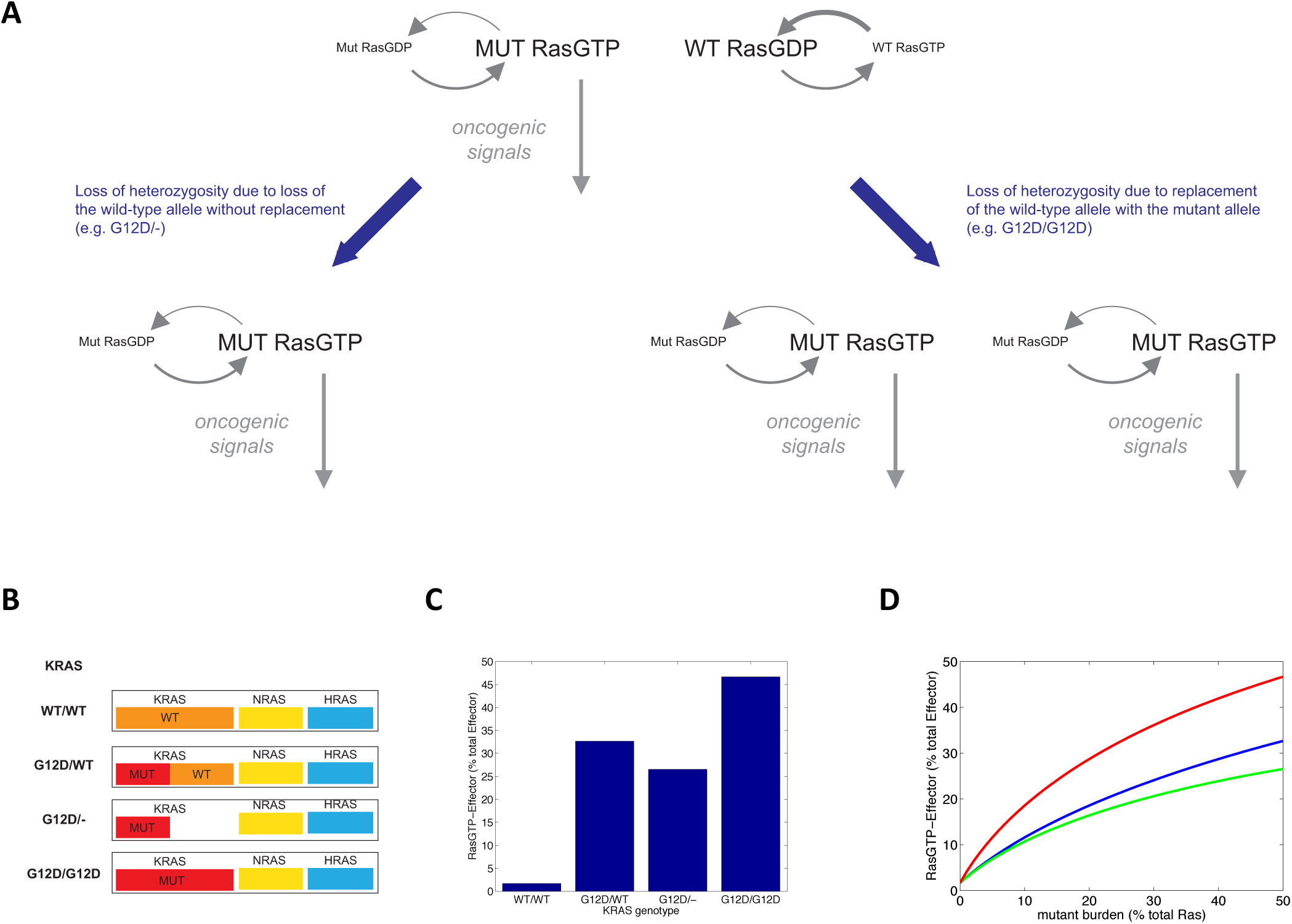
Analysis of RAS Loss of Heterozygosity as an argument for RAS tumor suppressor activity. A. Loss of heterozygosity that results in loss of wild-type RAS occurs when the wild-type allele is deleted and also when the wild-type allele has been replaced with the oncogenic allele. B. Schematic legend of the different modeled scenarios. C. RAS signal intensity for WT/WT, G12D/WT and the two LOH situations: G12D/- and G12D/G12D. D. RAS signal intensity for different mutant burdens of the G12D/WT case (blue), as well as the two situations for LOH: G12D/- (green) and G12D/G12D (red).

In our simulations, we find less RAS signal for G12D/- compared to G12D/WT conditions, and we find increased RAS signal for G12D/G12D (Figure 2C). If wild-type RAS counteracted the effects of oncogenic RAS, as is commonly hypothesized as part of the RAS TSG hypothesis, we would have expected to see more total RAS signal in both cases. Our simulations rather suggest that the quantity of RAS signal is sensitive to the quantity of oncogene present, and that loss of wild type via replacement of the wild-type allele with mutant causes higher signaling due to an increase in oncogene dosage. As the three RAS proteins KRAS, NRAS, and HRAS can be expressed within the same cell, 50% of the RAS in the cell will generally not be mutated, but rather a smaller fraction consistent with the relative expression of the mutant and non-mutant forms of RAS (Figure 2B). Performing these same simulations for different proportions of mutant RAS similarly finds that uncompensated loss of wild-type RAS always results in a decreased RAS signal intensity (Figure 2D). This suggests that the quantity of expressed oncogene expression is the critical factor driving RAS gene LOH.

Recent experiments explored the importance of NRAS mutant dosage using mouse genetic models. These experiments demonstrated no statistically significant evidence for RAS having TSG activity, but they did demonstrate strong evidence for dosage being important to oncogenic RAS signaling [20]. This study, other studies that investigated dosage [10], and our model all suggest that RAS LOH in cancer is better explained by the effects of increasing the expression of the oncogenic RAS mutant rather than by decreasing the expression of wild type RAS.

### Analysis of Argument #2: Some RAS effectors are tumor suppressors

RAS has many downstream effectors with which RAS specifically interacts in its GTP bound form. Among these are the established oncogenes BRAF and PI3K that are also commonly mutated in human cancer. There is also evidence that some RAS binding proteins, such as RASSF1, act as tumor suppressor genes [21]. The argument that RAS is a TSG often cites these putative TSG and argue that the TSG activity of RAS is through its downstream TSG partners (Figure 3A). If RAS is lost, the argument is that these downstream tumor suppressor functions are not activated. Some versions of this argument posit that wild-type RASGTP specifically signals through the tumor suppressor.

**Figure 3:**
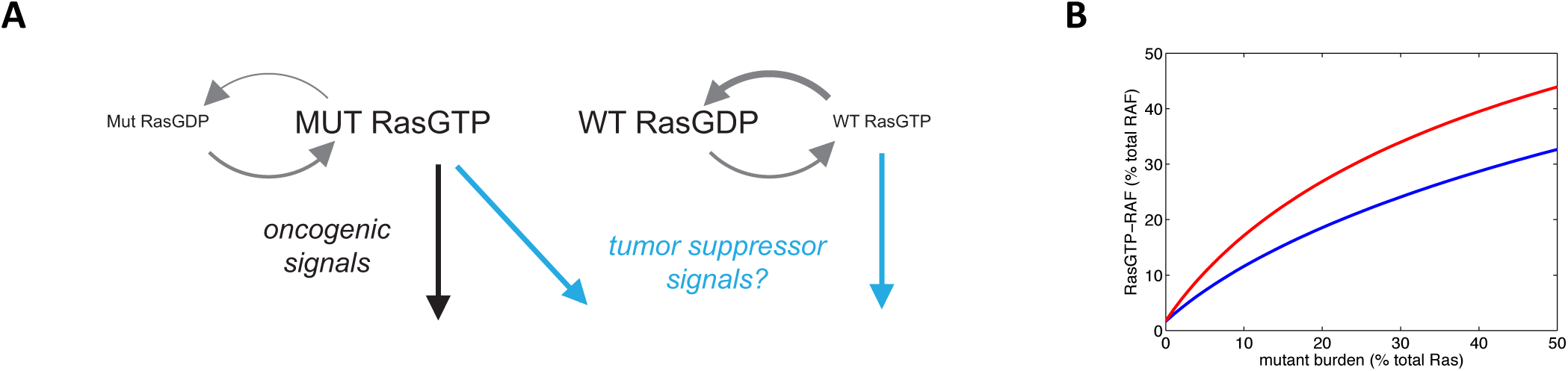
Analysis of the loss of putative tumor suppressors directly downstream from RAS. A. Schematic demonstrating hypothetical tumor suppressors that function downstream of RAS. B. Quantity of oncogenic RAS effector bound to RASGTP when the oncogenic and putative tumor suppressor effectors are both present (black), or after the tumor suppressor effectors are deleted (red).

In our original model, all RAS effectors are grouped into a single quantity of effectors. Here, the model was first extended to include two pools of RAS effectors (oncogenes, and “TSGs”). The total amount of effector was divided evenly between these two pools. Simulations were performed for both pools being present, and for the TSG pool being lost. The simulations demonstrate that loss of the “TSG” pool results in increased signaling through the oncogene pool (Figure 3B). It was not even necessary for the “TSG” pool to actually have tumor suppressor activity for their loss to result in increased oncogenic signaling, the loss of this pool of effectors simply results in increased signaling through the remaining oncogenic effectors. This simple competition for RAS binding may explain why loss of these RAS effectors appears to promote cancer; i.e. loss of expression for these putative tumor suppressors could actually result in increased oncogenic RAS signaling. Thus, a RAS effector may have a pattern of loss consistent with a tumor suppressor, but it’s loss is not selected for by changes in activity of downstream signaling partners but rather through a net decrease in competitive RAS binding reactions that can dilute active RAS-GTP away from its oncogenic effectors.

### Analysis of Argument #3: Dominant negative RAS inhibits oncogenic RAS signaling

Another argument used to argue that RAS has tumor suppressor activity is that dominant negative RAS can inhibit oncogenic RAS signaling. Dominant negative (DN) RAS are RAS mutant constructs that inhibit RAS signaling [22] (Figure 4A). These DN RAS mutants are predominantly GDP bound, and they bind to RAS GEFs with very high affinity. In this manner, DN RAS inhibits RAS GEF activity on the other RAS proteins in the same cell. DN RAS is usually used to inhibit WT RAS activation in experimental studies of physiological cell signaling. However, they have been used with “constitutively active” oncogenic RAS and have been found to diminish the phenotypes observed to be induced by oncogenic RAS [20, 23]. GEF activity is not widely thought of as being important to oncogenic RAS signaling [24]. With the assumption that oncogenic RAS is not dependent upon GEFs for activation, the DN data is therefore used to argue that RAS must have TSG activity [19]. The ‘RAS is a TSG’ argument posits that DN RAS should not block oncogenic RAS, so maybe elevated RASGDP from the DN RAS counteracts the elevated RASGTP from the oncogenic mutant. Such an argument assumes that there is a yet-to-be-discovered RASGDP signaling mechanism by which RAS exerts its TSG effects.

**Figure 4:**
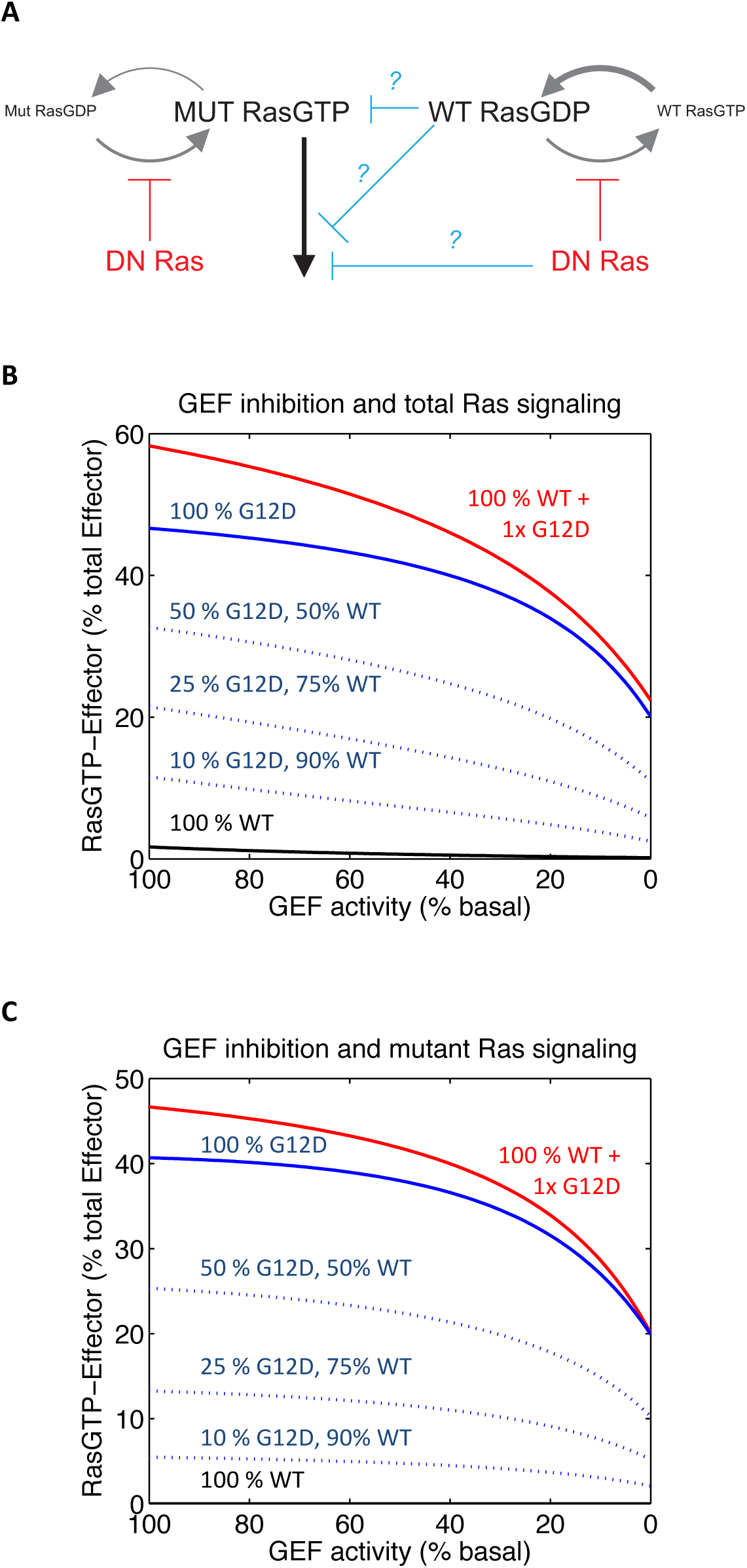
Analysis of dominant negative RAS on oncogenic RAS activation. A. Schematic demonstrating dominant negative (DN) RAS acts on RAS GEFs to block GTP loading. Tumor suppressor signals have been hypothesized to explain how DN RAS blocks oncogenic RAS signaling. A few of these hypothesized mechanisms by which DN RAS inhibits Mutant RASGTP signals are indicted in light blue. B. Dose responses for the G12D/WT network for different proportions of mutant RAS to WT RAS (blue) and for ectopic RAS G12D expression in addition to expression of endogenous WT RAS (red). The total amount of modeled ectopic expression is here modeled to be the same as the amount of endogenous RAS. C. The same dose responses as in B, but with only the mutant RAS bound to effector presented as the measure of RAS signaling.

In our model, we have previously demonstrated that inhibition of GEF activity within a network containing both oncogenic RAS and wild-type RAS can result in less total RAS signal [14]. We have found this modeling result to be one of the outputs that RAS biologists find least consistent with the conventional understanding of RAS biology, so we further elaborate upon this computational result here. Figure 4B presents dose responses of reduced GEF activity for networks with different proportions of wild type and oncogenic RAS G12D. We show that even when only oncogenic RASG12D is present, dominant negative RAS still inhibits levels of RASGTP. The dynamic equilibrium of mutant bound GDP and mutant bound GTP is, according to our model, influenced by the level of basally active GEF within the cell. Overall, our analysis finds that DN RAS inhibition of oncogenic RAS is actually consistent with available data on RAS regulation.

One might wonder in these cases whether DN RAS is blocking oncogenic RAS activation or wild-type RAS activation within the simulations, because oncogenic RAS can lead to wild-type RAS activation [14, 25-29]. The case of 100% of RAS being the G12D mutant clearly demonstrates that GEF activity is an important contribution to the activation of oncogenic mutants as more than half of total RAS signal is lost when the basal level of GEF activity is reduced to zero (Figure 4B). To further address this point, we limit our consideration of RAS signal to that generated by the mutant (Figure 4C). This clearly reveals decreases in mutant RAS signal. This further demonstrates that currently available biochemical knowledge of RAS signaling, as analyzed by our model, is consistent with DN RAS having the ability to inhibit mutant RAS via its known GEF inhibitory properties. This also reinforces the idea that RAS GEFs may serve as valuable drug targets due to their potential to reduce (but not abolish) signaling from oncogenic RAS mutants [30, 31].

The suggestion that dominant negative RAS can inhibit oncogenic RAS signaling has been met with some skepticism. Although the ability of DN RAS to inhibit signals from oncogenic RAS mutants has been demonstrated recently [20], some of the earliest studies using DN RAS found it to be quite ineffective at inhibiting oncogenic RAS [23]. Additionally, several recent studies have clearly demonstrated that oncogenic RAS activation remains dependent upon GEFs for full activation [32-34]. One potential variable that differed between the DN studies that might account for their different findings is the quantity of oncogenic mutant protein expressed. In the more recent study, the oncogenic NRAS mutant was expressed at physiological levels and in place of WT NRAS[20], whereas in the older studies oncogenic RAS was ectopically expressed in a cell that already had its full complement of RAS as WT RAS, which would result in supra-physiological levels of total RAS [23].

We performed additional simulations to evaluate whether differences in expression of wild-type and mutant RAS could explain the different, observed, responses to DN RAS. We find that supra-physiological expression of oncogenic RAS above and beyond the amount of endogenous RAS (all in the WT form) results in an overall higher level of total RAS signal and further shifts the dose response to the right (Figure 4B,C). This suggests that the early studies that indicated oncogenic RAS had very little sensitivity to DN RAS may have simply had too much total RAS and RAS signal to detect the effects of DN RAS.

### Analysis of Argument #4: The absence of a WT allele can amplify mutant RAS phenotypes

There are methods for chemically inducing cancer in mice. Several of these methods are known to induce RAS mutations in a very high proportion of the induced tumors [35]. One very strong argument for RAS having TSG activity came from studies that chemically induced tumors in wild-type mice with both alleles of *Kras* present (*Kras*+/+) and also in mice with one *Kras* allele deleted (*Kras*+/-) [36]. Intriguingly, a greater number of tumors were induced in mice with the *Kras*+/- genetic background. Cell culture experiments that compare Mutant/WT to Mutant/- conditions have found increased colony formation in soft agar, again suggesting that wild-type RAS may counteract the effects of mutant RAS [37, 38].

In Figure 2 we demonstrated that, according to our model, loss of wild type RAS in the presence of a RAS mutant should have less total RAS signal than a mutant accompanied by a wild-type allele. That the experimental data seems to contradict our model suggests that additional biological processes that are not included in our model may be at play. RAS having TSG activity is one possibility, but we have already demonstrated that the argument for RAS having TSG activity is less convincing than it first appears. We therefore considered an alternative hypothesis that that could explain these biological data.

When using our model, we have assumed that a loss of one copy of a gene would result in only half as much of that RAS protein being expressed. We hypothesize that RAS protein expression when only a single allele is present may be greater than half of the total RAS expression when both alleles are present. Supporting this hypothesis, previous measurements of Hras, Nras, and Kras in *Hras*^-^/^-^ mice found increased expression of Nras and Kras [39].

We used our model to determine the levels of RAS signal that would be found for various levels of compensated expression (Figure 5). We find that an increase of 10-30% from the remaining allele (which would result in 55%-65% as much RAS protein as would be observed for two alleles) is generally sufficient to cause a greater level of RAS signal. The previous experiments have not characterized RAS protein expression levels with such precision. The measurement of RAS protein expression within these genetic models is an important experiment to evaluate this alternative hypothesis, and should be included in future experiments.

**Figure 5:**
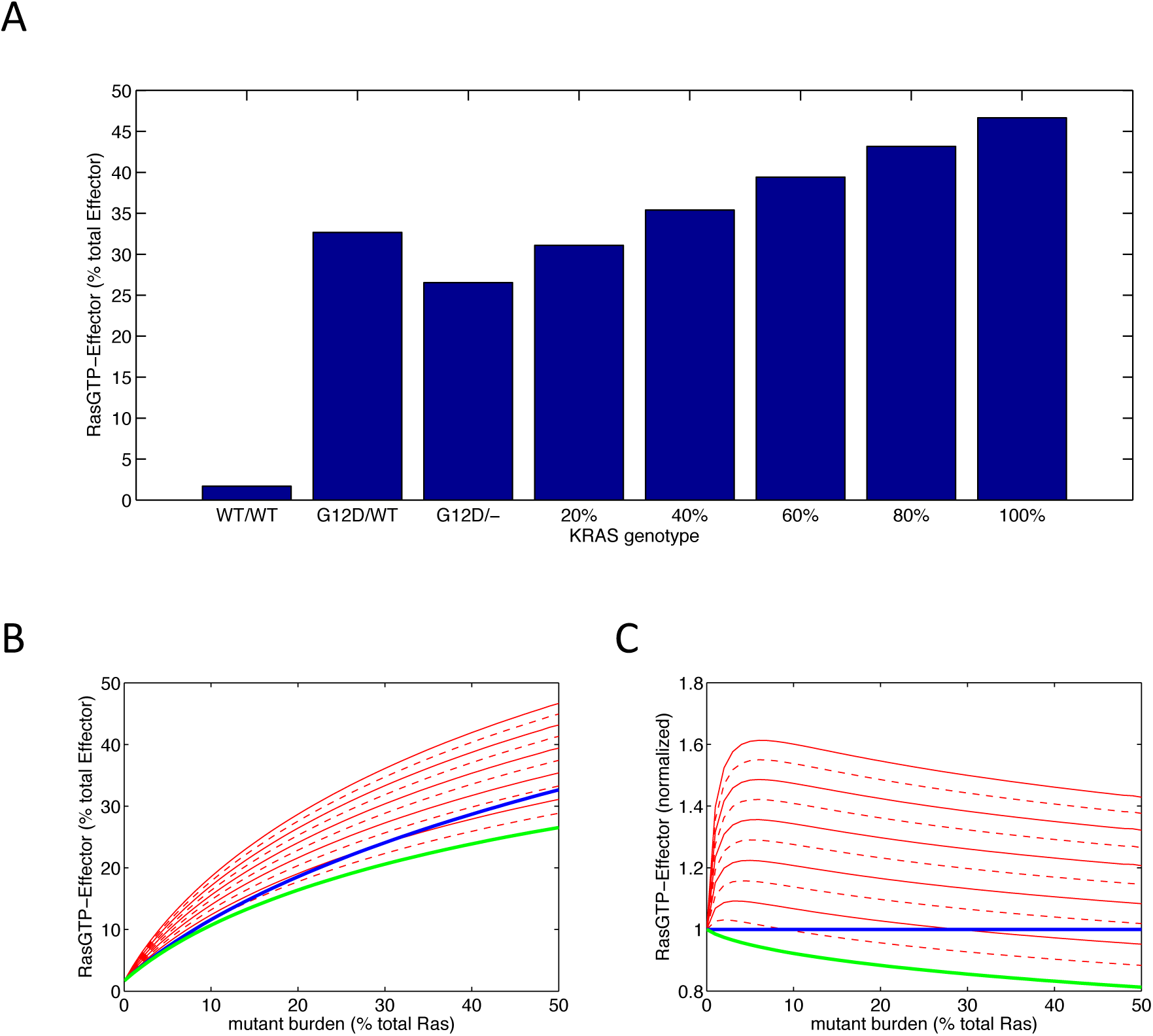
Analysis of RAS signaling in the presence of a null RAS allele. A. Levels of RAS signal for WT/WT, G12D/WT, and G12D/- conditions. We also consider various levels of increased expression from the remaining allele when one allele is lost. The percent of increased expression is indicated on the x-axis. B. G12D/WT (black), G12D/- with no compensation (green) and G12D/- with different levels of compensation portrayed in 10% increments; odd multiple levels are indicated with dotted red lines, even multiple levels are indicated with solid red lines. C. The data in B, normalized to the level of RAS signal found in the simulated G12D/WT conditions.

### Analysis of Argument #5: An additional WT RAS allele appears to counteract mutant RAS phenotypes

Recent experiments investigated the role of wild-type dosage on leukemia formation in mice that include one *Kras*G12D allele [40]. These experiments compared a *Kras* G12D/WT genotype with a *Kras* G12D/WT/WT genotype (i.e. a mouse with an additional WT *Kras* allele added). The experiments observed reduced leukemia development in the mice with the additional wild-type *Ras* allele [40].

In our model, an additional copy of wild-type RAS (e.g. *KRAS* G12D/WT/WT) should result in a slight elevation of RASGTP (Figure 6). This would go against our assumption that increased RASGTP results in a higher rate of cancer, and again could be interpreted as a sign that additional RAS biology that is not in our model must be at play. We again consider the hypothesis introduced in the analysis of argument #4, which was that there are mechanisms to control the total level of RAS protein expression. With an additional RAS allele, there may be cellular mechanisms to reduce expression from all three alleles. We modeled this possibility for different levels of compensation (Figure 6). At full compensation, there would be less oncogenic mutant expressed in the G12D/WT/WT context than in the G12D/WT context. In hypothesized conditions of partial compensation, our simulations suggest that 10-30% reduction in expression from each allele in the G12D/WT/WT condition is generally sufficient to reduce total RAS signal below that of the corresponding G12D/WT state. This suggests that experiments that measure the total level of G12D RAS in this system would be particularly valuable.

**Figure 6:**
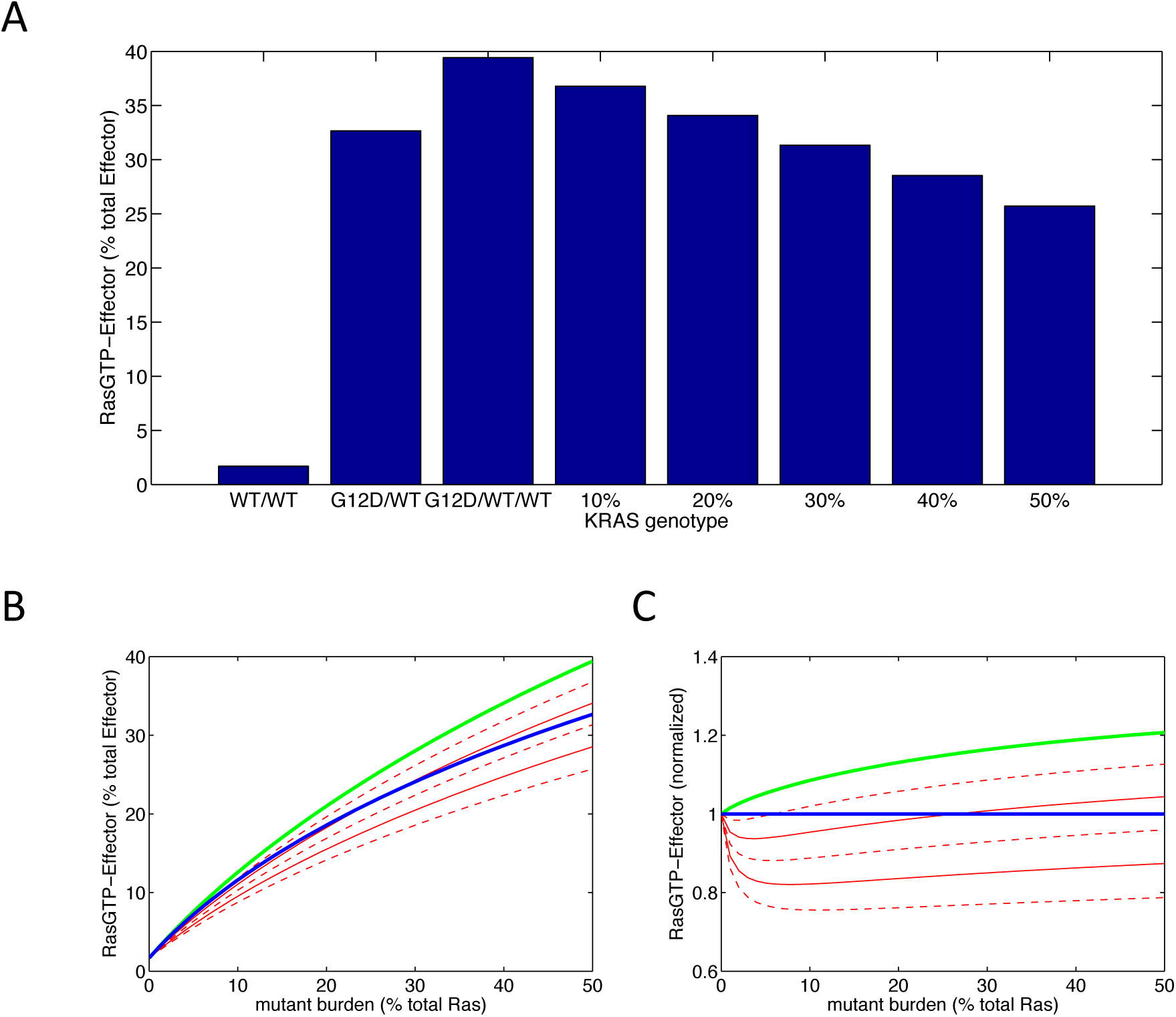
Analysis of RAS signaling in G12D/WT/WT conditions. A. Levels of RAS signal for G12D/WT/WT conditions assuming that the additional allele expresses at a level equivalent to the expression of each allele in G12D/WT cases, and then assuming that each of the three alleles are proportionally reduced by the percentage indicated. B. G12D/WT (black), G12D/WT/WT without compensation (green), and G12D/WT/WT with partial compensation (red lines) in 10% increments. Odd multiple levels are indicated with dotted red lines, even multiple levels are indicated with solid red lines. C. The data in B, normalized to the level of RAS signal found in the simulated G12D/WT conditions.

## DISCUSSION

We have computationally analyzed five separate types of experimental data that have previously been interpreted to support the argument that wild-type RAS has tumor suppressor activity. A central premise of these arguments is that known RAS biology cannot explain the data, but that RAS having tumor suppressor activity could. In our analysis, we find that three types of data believed to be inconsistent with RAS biology are actually consistent with known RAS biology. This suggests that the arguments that claim RAS must have TSG activity to explain existing experimental observations is much less strong than it has appeared.

To explain the other two types of data used to argue RAS has tumor suppressor activity, we introduced the assumption that RAS protein levels are subject to processes that limit production when additional copies of the RAS gene are present and that increase production when one RAS allele is absent. That RAS protein expression levels are not linear with gene dosage is arguably a simpler hypothesis than the tumor suppressor hypothesis, and it fits in well with the known mechanisms of RAS signaling. We highlight that relatively small changes in expression level may be sufficient to explain the previous experiments.

Recently published experiments offer new data that support our hypothesis of compensated expression from the mutant allele. This recent study used mouse genetic models of *Ras* mutants to study leukemia [41]. Of note, when they compared Ras expression between *Kras* G12D/WT and *Kras* G12D/null tissue, they detected a much less than 50% decrease in total Kras expression when the wild-type allele was deleted. Indeed, total *Kras* expression was barely changed when the wild-type allele was deleted. This suggests that there was increased expression from the *Kras* G12D allele. In further support of this interpretation, their measurements of Ras activation also found increased total KrasGTP when the wild-type allele was deleted. These new data are all consistent with our computational analysis and our hypothesis that increased expression from the mutant allele occurs when the wild-type allele is deleted.

Our computational analysis of the RAS as a TSG hypothesis demonstrates the potential value of using modeling to help evaluate experimental data. Biology would benefit from improved methods to determine whether unexpected data are indicative of new biological processes or are rather logically consistent with what is known but in a non-obvious way. Additionally, the use of modeling to evaluate data can help identify alternative, testable, hypotheses and important experimental controls. Wider incorporation of modeling into experimental biology may make biological progress more efficient.

One effort that may significantly advance the process of developing computational models of complex biological systems is the DARAPA “Big Mechanism” program. This effort aims develop computational technologies that can understand and model complex systems, and its initial focus is on RAS signaling in cancer [42, 43]. The program aims to develop computational methods that can read and interpret the biological literature and that can use the information gleaned from automated reading to develop models of the RAS signaling pathway. Our previous work demonstrates that available knowledge of RAS signaling can be used to develop a model with good predictive ability and that can uncover new biological behaviors that can then be experimentally confirmed [14, 16]. We believe the work presented here is also relevant to the “Big Mechanism” program. The RAS models to be developed as part of the “Big Mechanism” program will be based on an automated reading of the literature and the original authors’ interpretations of their data [43]. Our work here has investigated an area of RAS biology where different scientists have interpreted the same data differently. We also demonstrated behaviors that logically follow from what is known but may not be obvious to experts in the field. Author interpretations may represent one of the challenges that the Big Mechanism program will face when it develops computational models that are based upon authors’ interpretations of their studies.

## CONCLUSION

Our computational modeling find that the evidence for RAS having tumor suppressor activity is less strong than it has appeared. Many of the data that were assumed to be inconsistent with RAS biology are actually consistent with RAS biology. Data that were not consistent with RAS biology according to our model were consistent once an additional hypothesis of protein expression level not being linear with gene dosage was introduced. That RAS protein expression levels are not linear with gene dosage is a simpler hypothesis that fits in well with the known mechanisms of RAS signaling, whereas the RAS tumor suppressor gene hypothesis does not yet have a solid mechanistic foundation. Future efforts to argue that WT RAS has tumor suppressor activity should properly control for RAS dosage and should include quantitative measurements of RAS protein levels to rule out this plausible alternative hypothesis. Additionally, quantitative measurements of mutant protein expression in these model systems could experimentally evaluate our hypotheses about RAS protein expression.

